# Β_2_AR Agonists Sustain Thermogenesis and Leanness via *Sympathofacilitation*

**DOI:** 10.1101/2025.06.13.659468

**Authors:** Samson W. Cheung, David Sidarta-Oliveira, Lu Yao, Yitao Zhu, Gitalee Sarker, Sofia Lundh, Donald A. Morgan, Ji Qu, Sian Wilcox, Vladyslav Vyazovskiy, David J. Paterson, Dan Li, Kamal Rahmouni, Ana I. Domingos

**Author notes:** Corresponding (A.I.D.).

## Abstract

Human thermogenesis depends on β2-adrenoceptors (β2AR) expressed in thermogenic adipocytes, which are activated by norepinephrine released from sympathetic neurons. Whether β2AR also modulates thermogenesis via direct presynaptic action within sympathetic neurons has remained unclear. Here, we identify *Adrb2* expression in human and rodent cervical sympathetic neurons. β2AR agonism exerts neurotrophic effects and facilitates cholinergic responsiveness in mouse sympathetic neurons, indicating a *sympathofacilitatory* role. Selective deletion of β2AR in sympathetic neurons leads to impaired nerve activity in brown adipose tissue, sympathetic neuropathy, worsened fasting-induced hypothermia, and progressive obesity in chow-fed mice—without changes in food intake. These findings uncover a presynaptic role for β2AR in sustaining thermogenesis and regulating adiposity, suggesting *sympathofacilitation as* a therapeutic avenue for obesity.

**Highlights:**

- Human and mouse cervical sympathetic neurons express *adrenoceptor beta 2* (*Adrb2*)
- β2-adrenoceptor (β2AR) activation is neurotrophic and facilitates sympathetic neuronal excitability
- Loss of β2AR in sympathetic neurons leads to neuropathy in brown adipose tissue and reduced sympathetic activity
- Deletion of β2AR in sympathetic neurons exacerbates fasting-induced hypothermia and promotes obesity independently of food intake

## Introduction

Sympathetic neurons innervate white (WAT) and brown adipose tissues (BAT) to mobilize fat, dissipating stored energy in the form of heat.^1–3^ Sympathetic terminals release neuropeptide Y that sustains the biogenesis of thermogenic adipocytes^4^, and norepinephrine (NE) that triggers thermogenesis via β-adrenoceptor (βAR) signalling.^5^ βARs are the G protein-coupled receptors for NE and epinephrine (EPI), which can be divided into three subtypes, β1AR, β2AR and β3AR.^6^ Mouse thermogenesis is driven by β3AR but a recent study demonstrated β2AR as the predominant subtype in human brown adipocytes.^7^ High affinity β2AR agonists have been abused for weight loss and body building^8^ but they acutely activate BAT and increase energy expenditure in humans before muscle grows.^9–11^ In alignment with these findings, multiple genome-wide association studies (GWAS) revealed the association between *Adrb2* gene polymorphisms and obesity susceptibility, further corroborating the metabolic role of β2AR.^12–18^

Whereas postsynaptic roles of β2AR in adipocytes have been studied,^7^ a presynaptic role of β2AR in sympathetic neurons and its relevance for thermogenesis or fat mass control has never been directly tested. Here we unveil the autocrine role of β2AR in mediating *sympathofacilitation*^19^ that affects thermogenesis and body weight, independently of food intake.

## Results

### Sympathetic neurons express *Adrb2*

To confirm the expression of *Adrb2* in sympathetic neurons, we integrated 10 publicly available single-cell RNA-sequencing (scRNA-seq) datasets of sympathetic ganglia across anatomical levels and species. Total 166,099 cells from superior cervical (SCG), stellate and thoracic ganglia of mouse,^20–27^ rat^28^ and human^29^ datasets were integrated through an anchor-based approach with Seurat v3 canonical correlation analysis workflow, followed by TopOMetry dimensionality reduction (Figure 1A).^30,31^ 11,916 (7%) of cells were classified as neurons with pan-neuronal marker genes, *tubulin beta-3* (*Tubb3*) and *RNA binding fox-1 homolog 3* (*Rbfox3*) encoding neuronal nuclei (NeuN). Approximately 77% of neurons expressed *tyrosine hydroxylase* (*Th*), indicative of sympathetic signature, whereas 11% were double-positive for both *Th and Adrb2* (Figure 1B). All neurons were categorized into 13 distinct clusters shown in UMAP plot (Figure S1A). Interestingly, *Adrb1* and *Adrb3* were not detected in any cluster (Figure S1B). This highlights the importance of β2AR and raises the question of why *Adrb2* is preferentially expressed in sympathetic neurons. *In situ* hybridization assay with higher sensitivity that enables single mRNA molecule detection also confirmed the expression of *Adrb2* in mouse stellate ganglia, 26% of TH^+^ cells having *Adrb2* transcripts (Figure 1C).

**Figure 1.**
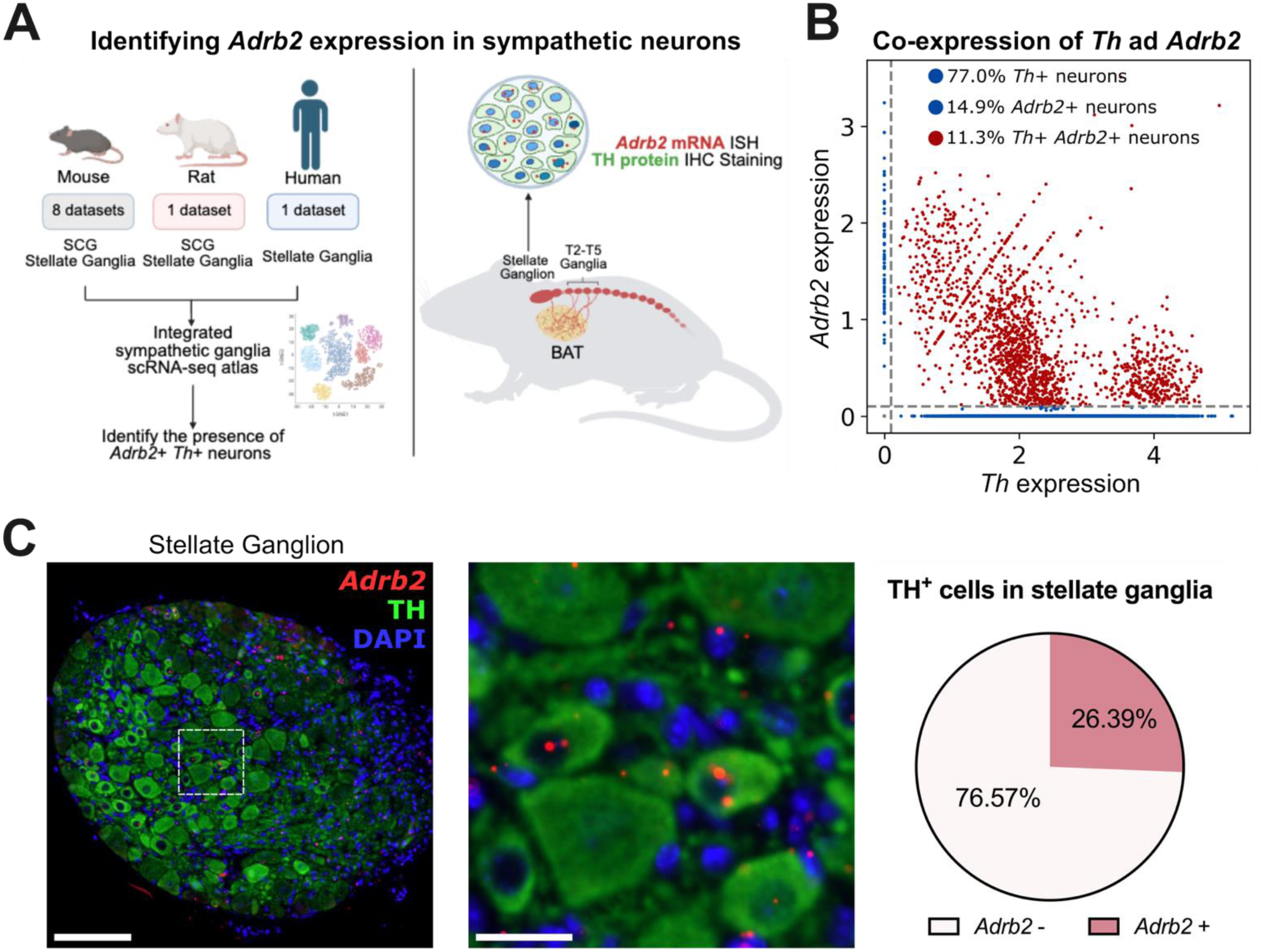
Sympathetic neurons express *Adrb2*. (A) Schematic diagram of identifying *Adrb2* in sympathetic nerves using integrated sympathetic ganglia scRNA-seq atlas, followed by *in situ* hybridization (ISH)-immunohistochemistry (IHC) staining. (B) Correlation plot of *tyrosine hydroxylase* (*Th*) and *Adrb2* gene expression in sympathetic ganglion neurons. (C) *In situ* hybridization of *Adrb2* mRNA with TH immunofluorescence staining in stellate ganglion Right: Percentage of TH^+^ cells with *Adrb2* mRNA (n = 4 mice). Scale bar, 100 μm for low magnification and 20 μm for zoom-in images.

### β2AR agonism facilitates sympathetic neuron activity and confers neurotrophic effects

Having identified β2AR in sympathetic neurons, we next investigated its functions using sympathetic neurons dissociated from mouse stellate ganglia (Figure 2A). We employed intracellular calcium ([Ca^2+^]_i_) imaging to assess the effect of β2AR on neuronal activity. Pharmacological activation by selective β2AR agonist clenbuterol (Clen) directly excited neurons and also led to greater [Ca^2+^]_i_ response to acetylcholine (ACh), a physiological sympathetic activator (Figures 2B-2D). The excitatory effects were abrogated by pre-treating neurons with butoxamine (But), a β2AR antagonist, proving that the Clen-evoked [Ca^2+^]_i_ response is dependent on β2AR activation (Figures 2B-2D). We then tested if β2AR plays a neurotrophic role in sympathetic neurons. We found that Clen treatment significantly induced neurite outgrowth of dissociated sympathetic neurons, whereas the effect was diminished in the presence of But (Figure 2E).

**Figure 2.**
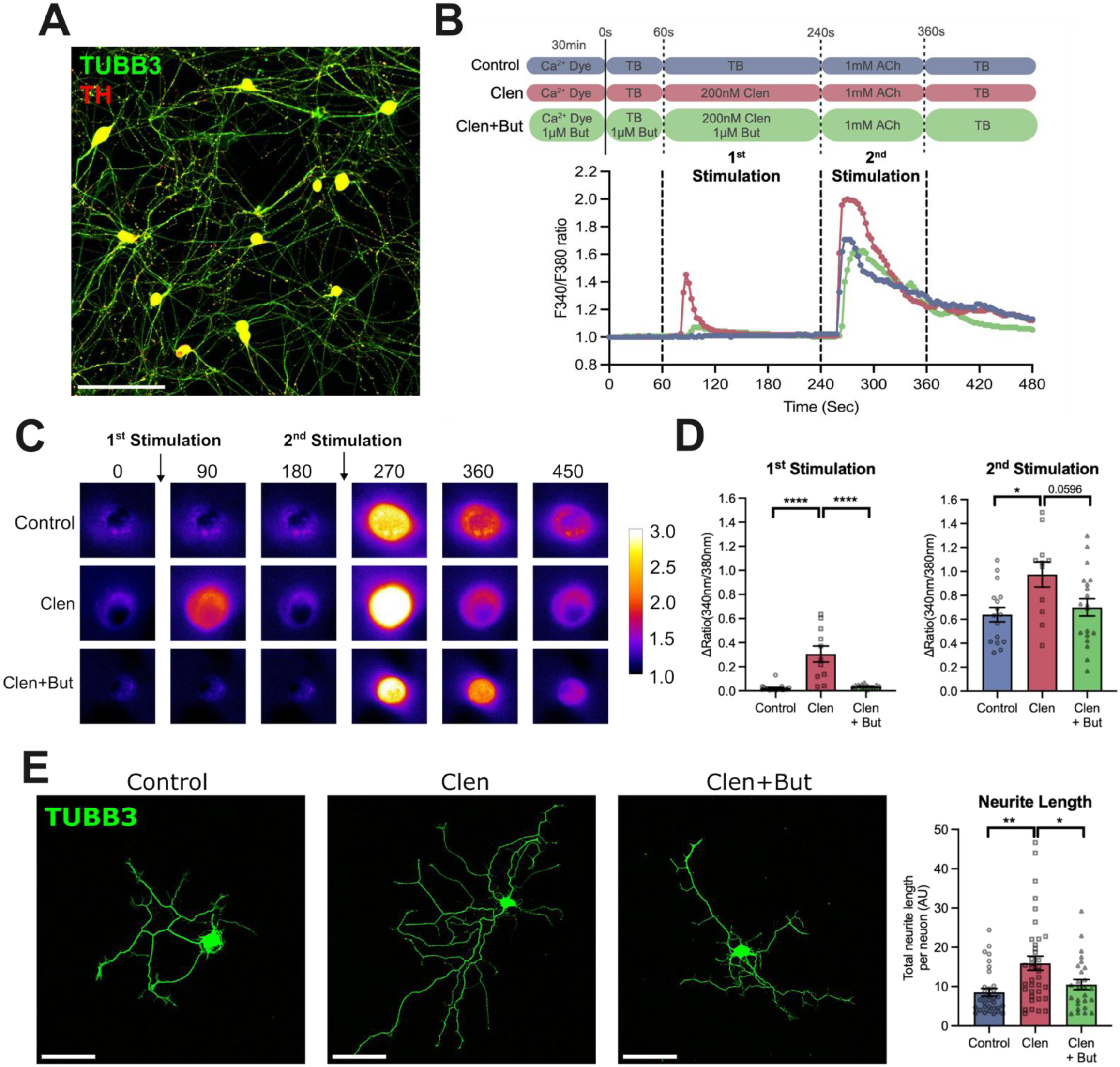
β2 agonist enhances sympathetic neuron activity and growth. (A) Primary sympathetic neurons from mouse stellate ganglia immuno-labeled for TUBB3 and TH. Scale bar, 100 µm. (B) Representative intracellular calcium ([Ca^2+^]_i_) response of sympathetic neurons treated with β2 agonist, clenbuterol (Clen) in the presence or absence of β2 blocker, butoxamine (But) (referring to 1^st^ stimulation), followed by acetylcholine (ACh) stimulation (2^nd^ stimulation). Tyrode’s buffer (TB) was used as basal wash solution. (C) Representative pseudocolor images of [Ca^2+^]_i_ changes in sympathetic neurons. (D) Amplitude of [Ca^2+^]_i_ response in the 1^st^ and 2^nd^ stimulation (n = 11-19 cells per group). (E) TUBB3 staining of mouse stellate sympathetic neurons treated with Clen in the presence or absence of But. Right: Quantification of neurite length (n = 26-37 cells per group). Scale bar, 100 µm Data are represented as mean ± SEM. Statistics: One-way ANOVA with Bonferroni multiple comparison test (D and E). *p < 0.05, **p < 0.01, ****p < 0.0001.

### Negligible number of cells in mouse brain co-expresses *Th* and *Adrb2*

To study the metabolic roles of sympathetic β2AR, we crossed *Th^Cre^* mice with *Adrb2^fl/fl^* mice to generate a knockout model with β2AR ablation in sympathetic neurons (herein named *Adrb2*^TH^KO). *In situ* hybridization assay confirmed that *Adrb2* was deleted in TH^+^ cells of *Adrb2*^TH^KO stellate ganglia (Figure S2). We also examined the specificity of *Adrb2*^TH^KO model by analysing public scRNA-seq dataset of the mouse brain,^32^ in which 2.3 million cells across regions were profiled. Only 0.015% of identified cells were found double-positive for *Adrb2* and *Th* (Figure S3A). Furthermore, hypothalamus, the major region that controls energy metabolism, showed negligible number of *Th^+^Adrb2^+^* cells (12 out of 162,869 hypothalamic cells) (Figure S3B). Altogether, *Th^Cre^*-mediated conditional *Adrb2* deletion is unlikely to cause off-target effects on the central nervous system that could confound the thermogenic phenotyping herein.

### β2AR sustains thermogenesis during fasting

BAT is a key metabolic organ controlled by sympathetic neurons.^33^ We then tested if sympathetic β2AR mediates BAT activity in response to metabolic challenges. Fasting is known to tone down the BAT sympathetic nerve activity (SNA) characterized by decreased NE turnover rate.^34^ Given the excitatory role of β2AR observed in our *in vitro* model, we hypothesized that β2AR sustains the basal BAT thermogenesis in the fasted state. *Adrb2*^TH^KO and *Adrb2^fl/fl^* littermate controls were challenged with 14-hour fasting (Figure 3A). *Adrb2*^TH^KO mice showed a significant reduction in both interscapular and rectal temperature after fasting relative to controls (Figures 3B-3D).

**Figure 3.**
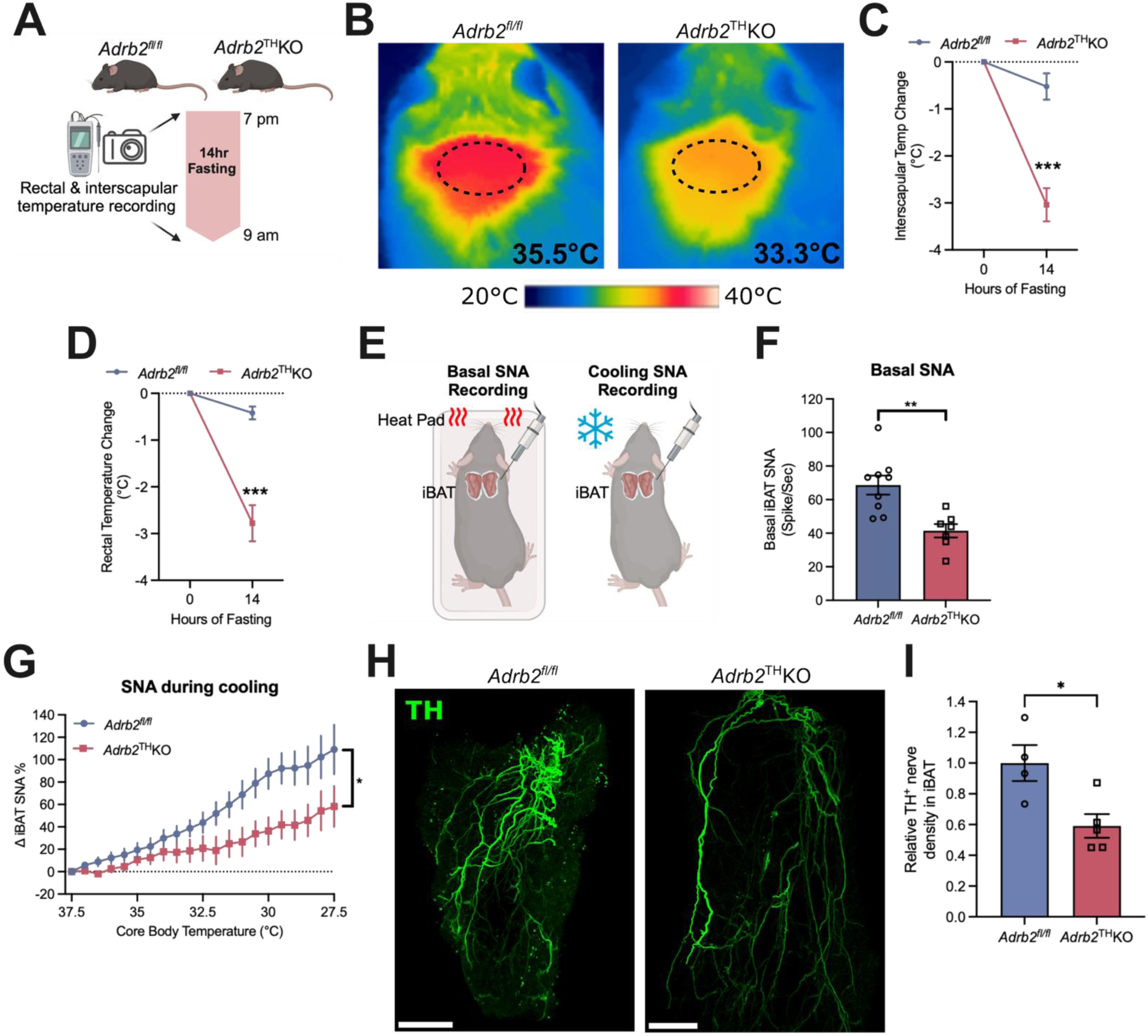
β2AR deletion in sympathetic neurons impairs brown adipose tissue thermogenesis and nerve activity. (A) Schematic depicting overnight fasting of *Adrb2^fl/fl^* and *Adrb2*^TH^KO male littermates. (B) Representative interscapular thermal images of mice before and after overnight fasting. (C-D) Interscapular region (C) and rectal (D) temperature changes after fasting challenge (n = 5-9 mice per group). (E) Schematic of basal and cooling-induced sympathetic nerve activity recording by electrophysiology (F) Basal sympathetic nerve activity (SNA) in iBAT of *Adrb2^fl/fl^* and *Adrb2*^TH^KO littermates (n = 7-9 mice per group). (G) iBAT SNA response to cooling (n = 7-9 mice per group). (H) Representative images of TH^+^ sympathetic nerve in iBAT. Scale bar, 1 mm. (I) Quantification of TH^+^ nerve fiber length normalized to total tissue volume in iBAT (n = 4-5 mice per group) Data are represented as mean ± SEM. Statistics: Unpaired two-tailed Student’s t-test (C-D, F and I); Two-way repeated-measures ANOVA (G). *p < 0.05, **p < 0.01, ***p < 0.001.

### β2AR is required for sympathetic activity and innervation in BAT

By employing *in vivo* electrophysiology on peripheral nerves entering the interscapular BAT (iBAT), we found that *Adrb2*^TH^KO had significantly lower iBAT SNA (Figures 3E and 3F). This indicates that presynaptic β2AR positively regulates iBAT SNA. To assess the role of sympathetic β2AR in cold-induced BAT thermogenesis, we subjected the anesthetized *Adrb2*^TH^KO mice and *Adrb2^fl/fl^* controls to cold by gradually lowering core body temperature from 37.5°C to 28.5°C. The increase in iBAT SNA triggered by cold was blunted in *Adrb2*^TH^KO mice (Figure 3G). These collectively show the necessity of sympathetic β2AR in iBAT activity during both fasting and cold challenges. In addition, we observed a significant reduction in TH^+^ nerve density of *Adrb2*^TH^KO iBAT, which aligns with the *in vitro* findings and supports the neurotrophic effect of sympathetic β2AR (Figure 3H).

### Loss of β2AR in sympathetic neurons causes adult-onset obesity independently of food intake

Finally, we examined the extent to which sympathetic β2AR influences body weight. We found that *Adrb2*^TH^KO mice on a chow diet gained more weight compared to *Adrb2^fl/fl^* littermate controls in early adulthood, and the trend attained statistical significance at 14 weeks of age (Figures 4A and 4B). This result was not confounded by stature as no difference in body length was seen (Figure 4C). Food intake remained unchanged relative to littermate controls (Figure 4D). Notably, *Adrb2*^TH^KO mice had larger and whitened iBAT as well as larger WAT pads (Figures 4E and 4F), whereas no change was seen in two key skeletal muscles, gastrocnemius and quadriceps (Figure 4G). These highlight the critical role of sympathetic β2AR in fat mass control.

**Figure 4.**
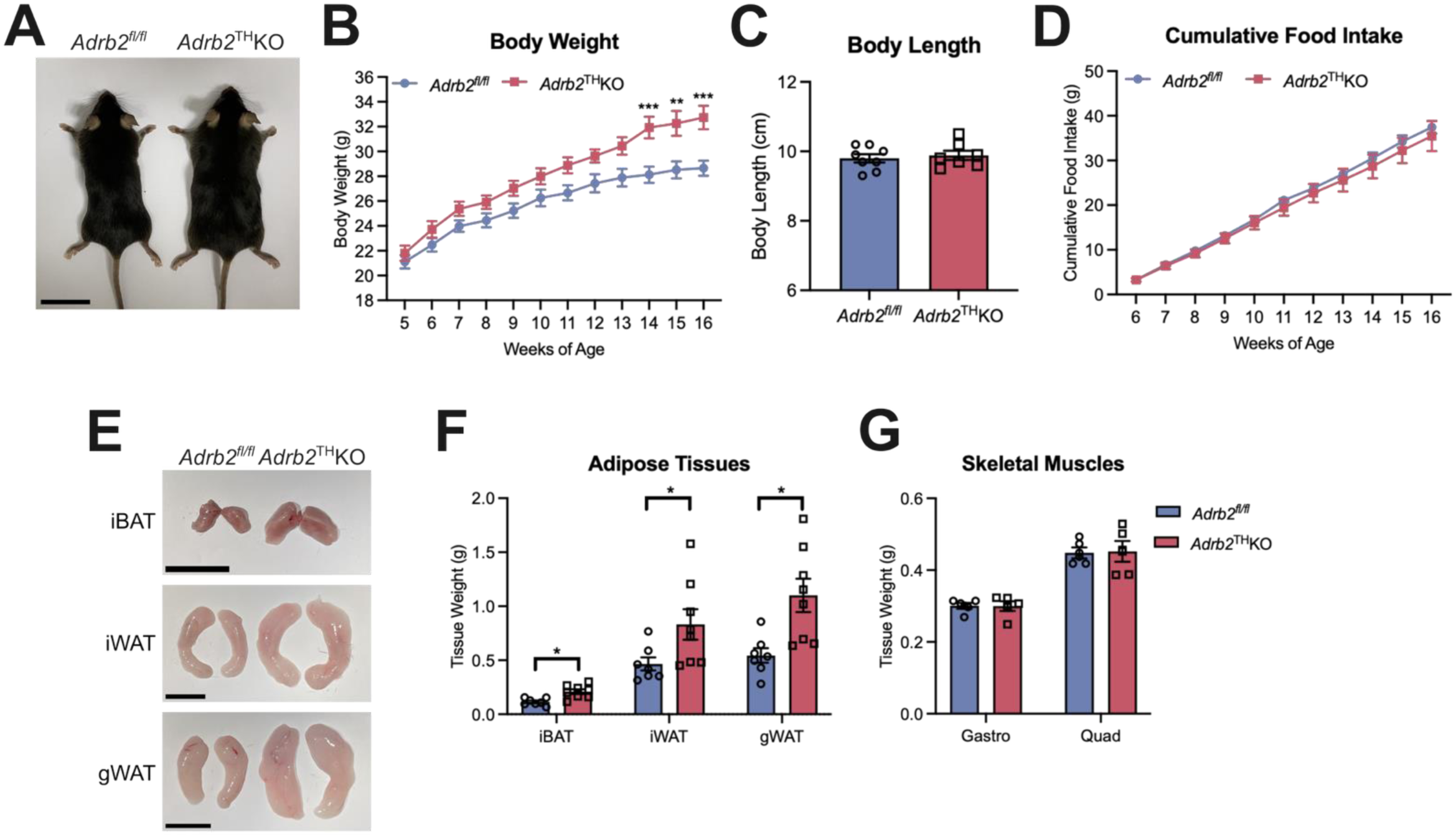
Loss of β2AR increases body weight and adipose tissue mass under standard chow diet. (A-B) Representative images (A) and body weight curve (B) of *Adrb2^fl/fl^* and *Adrb2*^TH^KO male littermates on standard chow diet (n = 10-12 mice per group). Scale bar, 3 cm. (C) Body length of mice at 16-17 weeks of age (n = 7-8 mice per group) (D) Cumulative food intake of mice (n = 4-5 cages per group) (E) Representative images of iBAT, gonadal (gWAT) and inguinal (iWAT) white adipose tissues. Scale bar, 1.5 cm. (F) Adipose tissue weight of mice at 16-17 weeks of age (n = 7-8 mice per group) (G) Gastrocnemius (Gastro) and quadriceps (Quad) weight of mice at 16-17 weeks of age (n = 5 mice per group) Data are represented as mean ± SEM. Statistics: Two-way repeated-measures ANOVA with Bonferroni multiple comparison test (B); Unpaired two-tailed Student’s t-test (F). *p < 0.05, **p < 0.01, ***p < 0.001.

## Discussion

Obesity is a modern epidemic with no clear explanation. Its increasing prevalence in the United States since 2000 cannot be linearly attributed to higher energy intake, as caloric consumption has remained unchanged over the same period.^35^ Instead, a decline in energy expenditure over the past three decades has been observed, primarily due to a reduced basal metabolic rate rather than changes in physical activity.^36^ Thermogenesis may be the underlying mechanism, as evidenced by three independent studies showing a gradual decline in core body temperature since the industrial revolution.^37^ Furthermore, BAT plays a critical role in cardiometabolic health, yet its biogenesis declines with both age and obesity.^38,39^ Thus, targeting BAT biogenesis and activation holds significant promise for therapeutic intervention for metabolic diseases.^38^

While β3AR signalling drives the thermogenic response in mouse adipocytes, human brown adipocytes are primarily activated through β2AR signalling.^7^ Specifically, β2AR agonism enhances glucose uptake in BAT and increases energy expenditure in young, lean individuals.^7,9–11^ Although β2AR agonists are thought to act directly on BAT, *in vivo*, a presynaptic effect in sympathetic neurons cannot be excluded. Notably, early pharmacological studies on isolated blood vessels suggested the existence of a β2AR-dependent presynaptic autocrine positive feedback loop in sympathetic nerves, facilitating NE release.^40–43^ However, this model has never been directly tested *in vivo*, nor has *Adrb2* expression been examined specifically in sympathetic neurons. Moreover, it was unknown whether this presynaptic β2AR-mediated autocrine loop impacts BAT thermogenesis, metabolism, or fat deposition.

To address this, we directly tested and confirmed *Adrb2* expression in sympathetic neurons of the stellate ganglion, which, in humans, innervates the mediastinum—where thermogenic BAT depots are located.^44^ In mice, the stellate ganglion innervates iBAT, where we performed electrophysiological recordings.^1^ Additionally, we uncovered a neurotrophic effect of β2AR agonism in adult sympathetic primary neurons, an effect previously observed only in embryonic forebrain cell lines.^45,46^ Consistent with our findings, the cAMP–protein kinase A (PKA) signalling pathway, known to support sympathetic neuron growth and survival, likely mediates this effect.^47,48^ Furthermore, we demonstrated that β2AR agonism has a *sympathofacilitatory* effect by enhancing sympathetic responses to acetylcholine *in vitro*.^19^ *In vivo*, *Adrb2* expression in sympathetic neurons is essential for maintaining both basal and cold-evoked sympathetic drive onto iBAT. Together, these findings provide direct evidence that a β2AR-mediated presynaptic autocrine positive feedback loop sustains sympathetic activity and thermogenesis.

Our results align with fundamental neuroscience principles, suggesting that, like most neurons, sympathetic outflow may be governed by activity-dependent neuronal plasticity and survival— a concept captured by Donald Hebb’s “use it or lose it” principle.^49^ A β2AR-dependent presynaptic autocrine feedback loop renders sympathetic neurons sensitive to their own activity, potentially explaining why cold exposure and exercise enhance sympathetic innervation in adipose tissue.^50,51^ The more a neural circuit is activated, the stronger it becomes; conversely, inactivity leads to weakening or loss. This principle may help explain how diet-induced obesity and leptin resistance, both of which suppress hypothalamic autonomic drive, lead to weakened sympathetic activity in adipose tissue—to the extent of promoting neuropathy.^52,53^ However, our findings suggest that presynaptic autocrine mechanisms, such as the β2AR-dependent loop, may also underly the “use it or lose it” principle to determine the density of sympathetic innervation of thermogenic fat. As the biogenesis of the latter is determined by sympathetic derived neuropeptide Y,^7^ then the “use it or lose it” neuronal principle may be mirrored on thermogenesis itself.

This model is further supported by our discovery that *Adrb2* loss-of-function in sympathetic neurons impairs thermogenesis during fasting and leads to adult-onset obesity despite normal food intake. In summary, we identify β2AR as a pivotal driver of sympathetic nerve activity and neurite plasticity, regulating both iBAT thermogenesis and adiposity. Our findings establish that the thermogenic effects of β2AR agonism are at least partially mediated through *Adrb2*+ sympathetic neurons, and it is plausible that this mechanism extends to humans.

## Experimental Model and Study Participant details

### Mouse Strains and Husbandry

Wild-type (WT) C57BL/6J mice were purchased from Charles River Laboratories. *Th*-specific *Adrb2*-deficient (*Adrb2*^TH^KO) mice were generated by crossing *Adrb2^fl/fl^* as previously described^54^ with *Th^Cre^* mice from Jackson Laboratory (Jax #008601). All mice used were on the C57BL/6J background. Mice were bred and maintained at the University of Oxford Animal Facility under specific pathogen free conditions, with constant temperature (21±1°C) and humidity (50±10%) under a 12-hour light/dark cycle (light period 7:00 am – 7:00 pm). Food and water were supplied *ad libitum* unless mentioned otherwise. Male mice aged 8-16 weeks were utilized for metabolic phenotyping as described in the corresponding figure legends. All experiment procedures were performed in accordance with the University of Oxford Institutional and UK Home Office regulations, and the University of Iowa Animal Care and Use Committee.

### Primary Cell Culture

Stellate ganglia were dissected from WT mice aged 4-5 weeks and cleaned of all surrounding connective tissues before digestion with 1 mg/mL collagenase type IV (Gibco 17104-019) in Hank’s balanced salt solution (Gibco 14025100) for 30 min at 37°C. Subsequently, ganglia were incubated by 0.25% trypsin (Gibco 25050014) for 15 min at 37°C, followed by trituration with 23-, 25- and 27-gauge needles to yield single-cell suspensions. Cells were seeded on glass bottom chamber slides (Ibidi 80807) or Nunclon cell culture plate (ThermoScientific 176740) coated with poly-D-lysine (Sigma P0899) and laminin (Roche 11243217001), and cultured in Leibovitz’s L-15 medium (Sigma L5520) supplemented with 10% fetal bovine serum (Sigma F9665), 24 mM sodium bicarbonate (Sigma S5761), 38 mM D-(+)-glucose (Sigma G8270), 0.5% penicillin-streptomycin (Gibco 15140) and 100 ng/mL nerve growth factor-7S (Sigma N0513)

## Method Details

### Dual *in situ* hybridization-immunohistochemistry (ISH-IHC) staining

Mice were euthanized with an overdose of pentobarbital and perfused transcardially with PBS. Stellate ganglia were fixed with 10% neutral buffered formalin (Sigma) overnight at room temperature and subsequently transferred to 70% ethanol. Samples were dehydrated, embedded in paraffin, sectioned at a thickness of 5 μm and mounted onto SuperFrost Plus slides (Fisher Scientific). Dual ISH-IHC was performed using RNAscope LS Multiplex Fluorescent Reagent Kit (ACD 322800), RNA-Protein Co-detection Ancillary Kit (ACD 323180) and BOND-RX Research Stainer (Leica Biosystems) according to manufacturer’s protocol. Tissue sections were co-stained for *Adrb2* mRNA and TH protein using RNAscope *Adrb2* probe (Mm484 Adrb2-XRn-C2, ACD 1172758-C2) and primary anti-TH antibody (1:1000, Millipore AB152), respectively. Probes targeting *Th* mRNA (Mm-Th-XRn-C2, ACD 870118-C2) and *Ppib* mRNA (Mm-Ppib, ACD 313918-C2) were used as positive ISH controls, while negative control was performed using DapB probe (ACD 312038-C2). As a positive control for IHC, primary anti-GLUT1 antibody (1:200, ThermoFisher PA5-16793) was used, and as negative control for IHC signal, an IHC protocol omitting primary antibody was used. Primary antibodies were detected with a Brightvision goat anti-rabbit poly-HRP (Immunologic) followed by Opal 690 Fluorophore Reagent (1:500, Akoya Bioscience FP1497001KT). ISH signals were visualized using iFlour 546 Styramide (1:500, AAT Bioquest/VWR 45025). Tissue slides were counterstained with DAPI before mounting with anti-fade medium (ThermoFisher Scientific). Images were acquired using Olympus VS200 slide scanner using a 20x (0.8 NA) air objective and DAPI/FITC/CY3/CY5 filter set. The number of TH+ *Adrb2*+ cells in the whole cross-sectional area was calculated by Image J with Cell Counter plugin. Approximately 200-400 intact TH-positive cells with clear cellular boundaries and stained nucleus were identified in each ganglion sample.

### Intracellular calcium imaging

Primary sympathetic neurons were dissociated from stellate ganglia, and seeded on poly-D-lysine- and laminin-coated coverslips as described above. Neurons were then cultured for 3 days and medium was changed every 24 hours. Following this, neurons were loaded with 5 µM Fura2, AM (Invitrogen F1221) in Tyrode’s buffer (135 mM NaCl, 4.5 mM KCl, 20 mM HEPES, 11 mM glucose, 1 mM MgCl2 and 2 mM CaCl2) for 30 min at 37°C. 1 µM Butoxamine, But (Santa Cruz SC-234233) was also loaded for the corresponding experiment group. Coverslip was then transferred to a temperature-controlled (37°C) chamber superfused with Tyrode’s buffer at a flow rate of 2 mL/min. Neurons were visualized using Nikon Diaphot microscope with 40X oil immersion lens under brightfield illumination. Neurons with round shaped cell bodies and neurites that were longer than the diameter of cell bodies were selected for calcium imaging assay.

Calcium levels at the cell bodies of neurons were recorded by exciting alternately at 340 and 380 nm at interval of 3s and the emission wavelength was set to 510 nm. During the experiment, the basal response was recorded for 1 min before stimulation of 200 nM clenbuterol, Clen (Sigma C5423) with or without 1 µM But in Tyrode’s buffer for 3 min. For control group, no Clen or But was administered. Subsequently, neurons were stimulated with 1 mM acetylcholine, ACh (Sigma, A2661) for 2 min before washing with Tyrode’s buffer. Calcium response was expressed as the ratio of fluorescence 510 nm emission on excitation 340 nm and 380 nm (F340/F380) or the normalized ratio (ΔF=F_1_-F_0_; F_1_ is the peak F340/F380 after Clen/ACh stimulation; F_0_ is the average of F340/F380 taken 15 seconds before stimulation). Data was recorded and processed using MetaFluor (Molecular Devices).

### Neurite growth assay and imaging

Primary sympathetic neurons were isolated and plated on glass bottom chamber slides (Ibidi 80807) as described above. PBS, 200 nM Clen and/or 1 µM But were added to medium 6 hours after plating. Following 48 hours of culture, neurons were fixed with 4% paraformaldehyde (PFA), followed by incubation with blocking/permeabilization buffer containing 3% bovine serum albumin (Sigma), 2% goat serum (MP Biomedicals), 1% Triton X-100 (Fisher Bioreagent) for 1 hour. Cells were then incubated with primary anti-TH (1:500, Millipore AB152) and anti-TUBB3 antibodies (1:500, BioLengend 801202) overnight at 4°C. Following washes with PBS, cells were incubated with DAPI (1:1000, Invitrogen), Alexa-488 (1:500, Invitrogen) and Alexa-546 (1:500, Invitrogen) conjugated secondary antibodies for 1 hour at room temperature. Samples were then washed with PBS before mounting with anti-fade medium (Invitrogen). All TH^+^ TUBB3^+^ cells with intact cell bodies and neurites were imaged using LSM880 confocal microscope and neurite length were quantitated by Image J with Neuron J plugin^55^. 6-9 replicate wells were included for each experimental condition.

### Fasting and temperature recording

Male mice at 8 to 9 weeks of age were singly housed and the interscapular region was shaved at least 2 days before any treatment performed. Thermal images of interscapular region were captured before and after 14-hour fasting (from 7 pm to 9 am) by E96 Advanced Thermal Imaging Camera (FLIR) as described previously.^29^ Mice were moving freely in the cage without any restraint during thermal imaging. Average interscapular surface temperature was calculated from 20-30 images of each mouse and timepoint using Thermal Studio Suite software (FLIR). Rectal temperature was measured using thermal probe (Physitemp).

### Single-cell RNA sequencing analysis

To create the integrated sympathetic ganglia scRNA-seq atlas, 10 public datasets comprising single-cell and single-nucleus RNA-seq of sympathetic ganglia were integrated using Seurat v3 canonical correlation analysis workflow, following standard quality control and log-normalization.^30^ Downstream clustering and visualization were performed with TopOMetry.^31^ Co-expression analysis of *Th* and *Adrb2* was performed with an expression threshold of 0.5. Information including the species, sequencing methods and ganglion samples used in each study has been summarized in Supplementary Table 1.

For analysis of mouse brain data, a publicly available whole-brain atlas was retrieved,^32^ and total 2,341,350 cells were used for log-normalization and querying the expression of *Th* and *Adrb2*. All code used for the aforementioned analyses will be available in a public repository.

### *In vivo* electrophysiology recordings

Male mice at 8 to 9 weeks of age were anesthetized with ketamine (91 mg/kg) and xylazine (9.1 mg/kg). To sustain anaesthesia throughout experiment, a micro-renathane tubing (MRE-40, Braintree Scientific) was inserted into the right jugular vein for infusion of α-chloralose (initial dose of 12 mg/kg and sustaining dose of 6 mg/kg per hour). The left carotid artery was cannulated with another MRE-40 catheter for monitoring heart rate and arterial pressure. Body temperature was also continuously monitored with rectal probe and kept constant at 37.5°C using temperature controller with heating pad and lamp (Physitemp Model MCAT2).

To access the nerve bundle innervating iBAT, the mouse was positioned on its ventral side and an incision in the nape of the neck was performed. A bipolar platinum-iridium electrode (A-M Systems) attached to high-impedance probe (Grass Instruments, HIP-511) was positioned under the nerve and covered with silicone gel (World Precision Instruments). The nerve signal was amplified by 10^5^ times, and filtered with 100-Hz and 1,000-Hz cutoffs with the nerve traffic analysis system (University of Iowa Bioengineering Model 706C). Following this, signal was transferred to speaker system and oscilloscope (Hewlett-Packard Model 54501A) to monitor the audio and visual quality of SNA recordings. Signal was directed to MacLab analogue-digital converter (AD Instruments Castle Hill Model 8S) containing software (MacLab Chart Pro Version 7.0) for analysing the number of spikes/second beyond the background noise threshold. Baseline iBAT SNA was continuously recorded for 30 min. Following this, core body temperature of mouse was lowered from 37.5°C to 27.5°C (approximately 0.25°C decrease per min) by removing all heat sources and placing ice packs on the metal platform. iBAT SNA was continuously measured during the entire cooling challenge. By the end of recording, the mouse was euthanized and background noise was determined to normalize iBAT SNA measurements.

### Whole-mount immunofluorescence staining

iBAT samples were delipidated, immunolabeled and cleared using modified iDISCO method.^4,56^ All incubation steps were performed with gentle shaking. Briefly, mice were euthanized with an overdose of pentobarbital, followed by intracardiac perfusion with PBS and then 4% PFA. Samples were post-fixed with 4% PFA overnight before dehydration in a graded series of 20%, 40%, 60%, 80%, and 100% methanol for 30 min each. Samples were incubated with 66% dichloromethane (DCM) (Sigma 270997) in methanol overnight. After washing with 100% methanol for 1 hour twice, samples were blenched using 5% H2O2/methanol overnight at 4°C and rehydrated in 80%, 60%, 40% and 20% methanol for 30 min each. Subsequently, samples were incubated in permeabilization buffer (20% DMSO, 0.2% Triton X-100, 0.3 M glycine in PBS) at 37°C for 48 hours and blocked in blocking buffer (10% DMSO, 0.2% Triton X-100, 0.02% sodium azide and 5% goat serum in PBS) for 48 hours.

Samples were incubated in anti-TH antibody (1:500, Millipore AB152) in dilution buffer (5% DMSO, 0.2% Tween-20, 10 ug/mL heparin, 0.02% sodium azide and 3% goat serum in PBS) at 37°C for 9 days, and then washed with PTwH buffer (0.2% Tween-20, 10 ug/mL heparin, and 0.02% sodium azide in PBS) for 6 times, 1 hour each. Following this, samples were incubated with Alexa-647 conjugated secondary antibody (1:500, Invitrogen A21244) in dilution buffer (0.2% Tween-20, 10 ug/mL heparin, 0.02% sodium azide and 3% goat serum in PBS) at 37°C for 7 days. After washing with PTwH buffer for 6 times, immunolabelled tissues were embedded in 1% agarose, dehydrated in 20%, 40%, 60%, 80% and 100% methanol for 1 hour each, and finally 100% methanol overnight at room temperature. Embedded samples were delipidated in 100% DCM for 2 hours twice before overnight clearing with dibenzyl ether (DBE) (Sigma 108014). Samples were then washed with ethyl cinnamate (ECi) (Sigma 112372) for 2 hours and then stored in fresh ECi in dark at room temperature.

Cleared samples were imaged on Miltenyi UltraMicroscope Blaze or Ultramicroscope II. 3D reconstruction was performed using Imaris 10 software (Bitplane) with volume rendering function. Orthogonal projection was utilized for representative 3D images shown in figures. The volume (µm^3^) and neurite length (µm) of iBAT were calculated by Imaris Surface FilamentTracer tools respectively. Sympathetic nerve density was represented as the ratio of total neurite length by tissue volume.

### Statistical analysis

All data are expressed as Mean ± SEM. All statistical analyses were performed with GraphPad Software 10 (GraphPad). p < 0.05 was considered statistically significant. Statistical test and sample size are indicated in the corresponding figure legends.

## Acknowledgments

We thank the University of Oxford Biomedical Services and Dunn School Bioimaging Facility for their help on this project. This work has been supported by grants from Wellcome Trust– Howard Hughes Medical Institute International Research Scholar Award (208576/Z/17/Z), ERC consolidator award (ERC-2017 COG 771431), 2022 Pfizer ASPIRE Obesity award (70591281), NIH PTE (5UM1DK105554-09) sub-award (5000826-5500002717), BBSRC (BB/Y006488/1), University of Oxford Croucher DPhil Scholarship awarded to S.W.C., Novo Nordisk Postdoctoral Fellowship awarded to G.S., Shineroad Industry Developments Co. Ltd. Scholarship awarded to Y.Z and China Scholarship Council Fellowship awarded to L.Y. K.R. is supported by NIH (R01 HL162773 and R01 HL172944), VA (I01 BX004249 and IK6 BX006040) and the University of Iowa Fraternal Order of Eagles Diabetes Research Center. S.W. is supported by NC3Rs (NC/S001689/1).

## Author Contributions

A.I.D. conceptualized the study. D.S.-O. constructed the integrated sympathetic ganglia scRNA-seq atlas and analyzed mouse brain scRNA-seq data. G.S. established sympathetic ganglia ISH-IHC protocol with S. L. and assisted S.W.C with stellate ganglia dissection, processing and image analysis. All experiments of sympathetic neuron culture were designed and performed by S.W.C. and J.Q. D.L. made intellectual contributions to sympathetic neuron isolation and activity measurement. S.W.C. and L.Y. did weight acquisition and fasting challenge of *Adrb2*^TH^KO mice. V.V., S.W. and S.W.C contributed to *in vivo* thermal imaging experiments and helped discover hypothermic phenotype of fasted *Adrb2*^TH^KO mice. Y.Z. provided advice on metabolic phenotyping and established the protocol of thermal imaging. D.A.M. and K.R. designed and performed *in vivo* electrophysiology. S.W.C. and A.I.D. wrote the manuscript with input from all authors.

## Key resources table

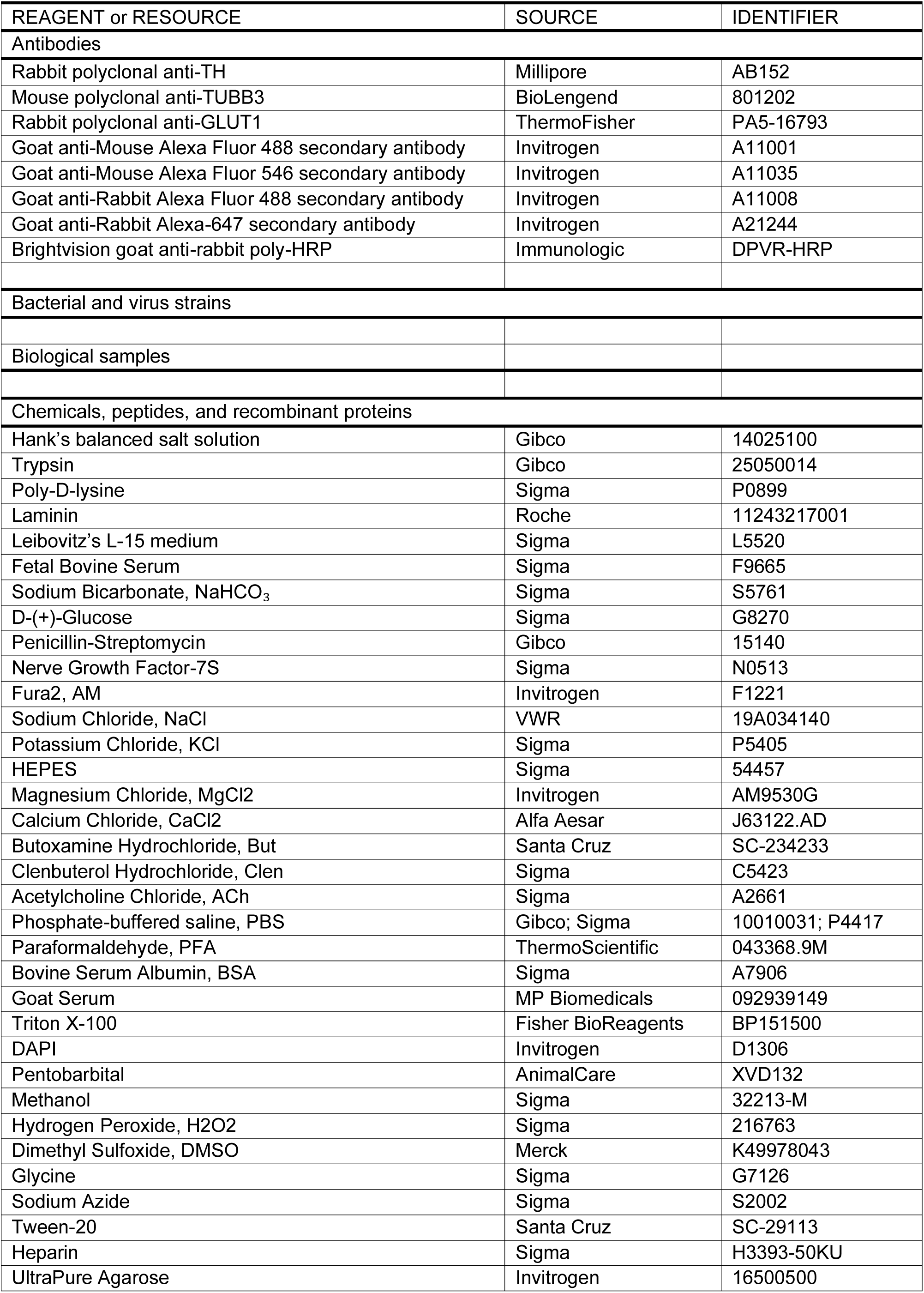
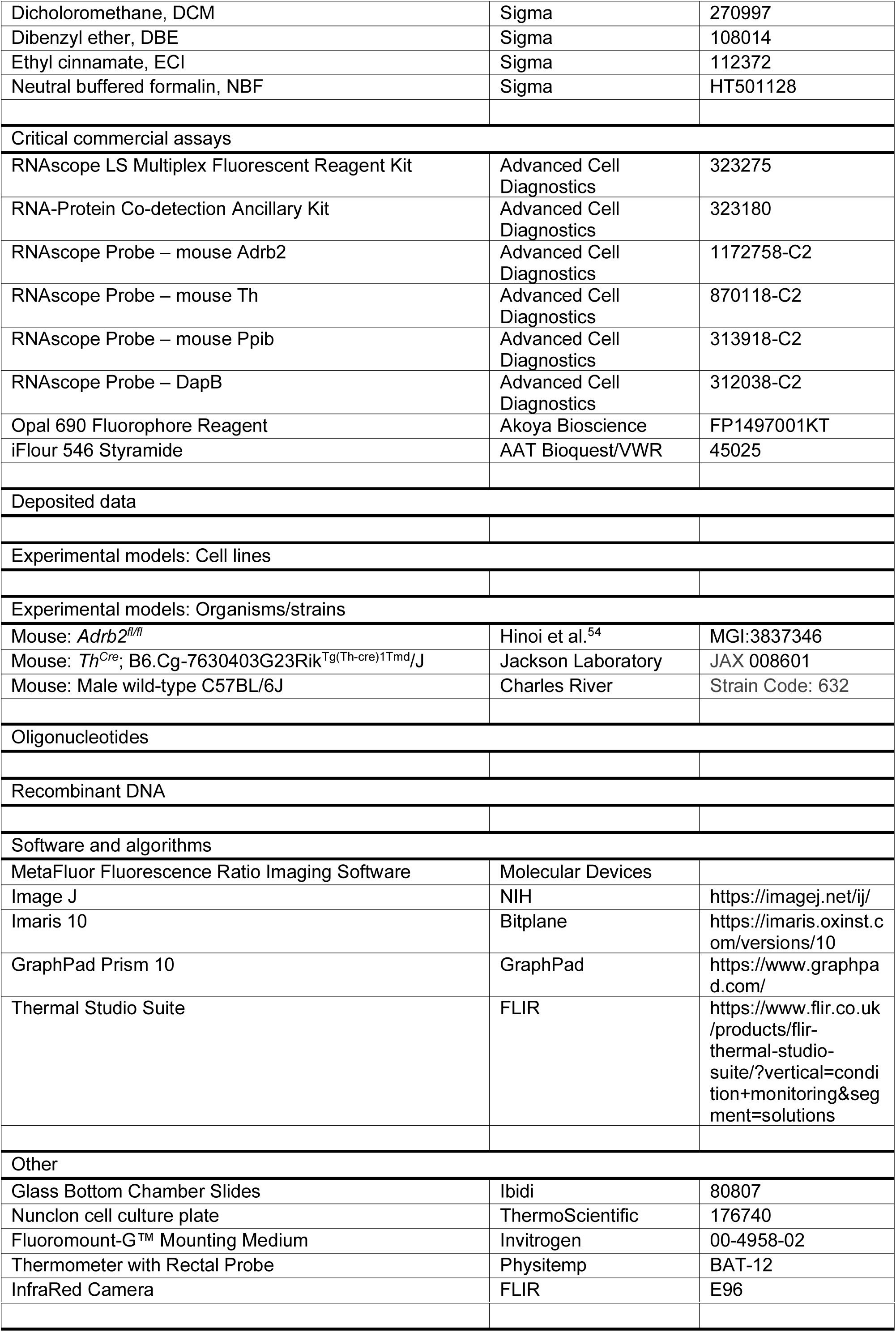

---

**Supplementary Figure 1.**
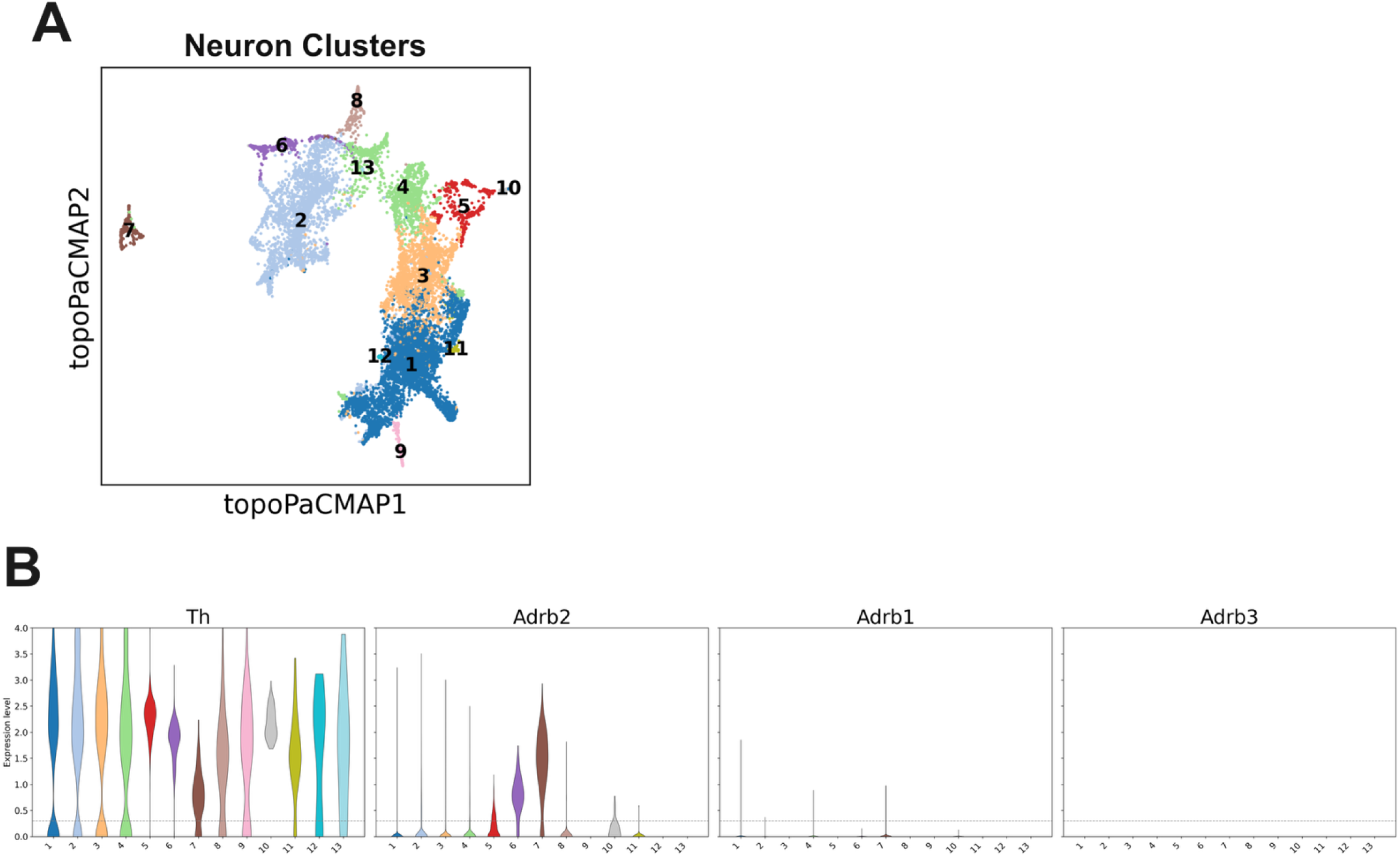
Sympathetic neurons do not express *Adrb1* and *Adrb3*. (A) UMAP plot of integrated sympathetic ganglia scRNA-seq atlas showing 13 neuron clusters. (B) Violin plot indicating *Th, Adrb2, Adrb1* and *Adrb3* expression levels among neuron clusters.

**Supplementary Figure 2.**
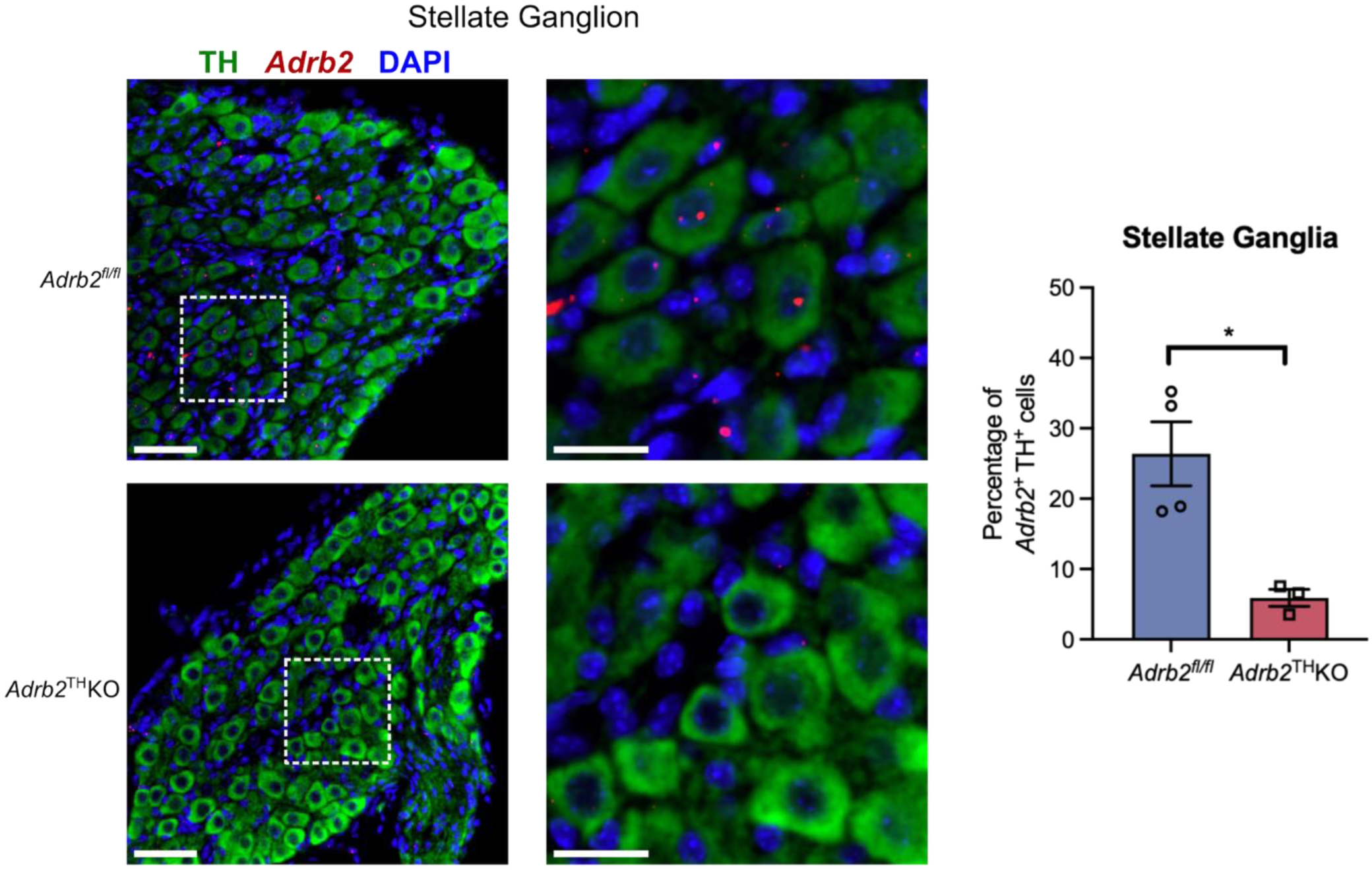
*Adrb2* is ablated in the TH^+^ sympathetic neurons of *Adrb2*^TH^KO stellate ganglia. *In situ* hybridization of *Adrb2* mRNA with TH immunofluorescence staining in *Adrb2^fl/fl^* and *Adrb2*^TH^KO stellate ganglion. Right: Percentage of TH^+^ cells with *Adrb2* mRNA (n=3-4 mice per group). Scale bar, 50 μm for low magnification and 20 μm for zoom-in images. Data are represented as mean ± SEM. Statistics: Unpaired two-tailed Student’s t-test. *p < 0.05.

**Supplementary Figure 3.**
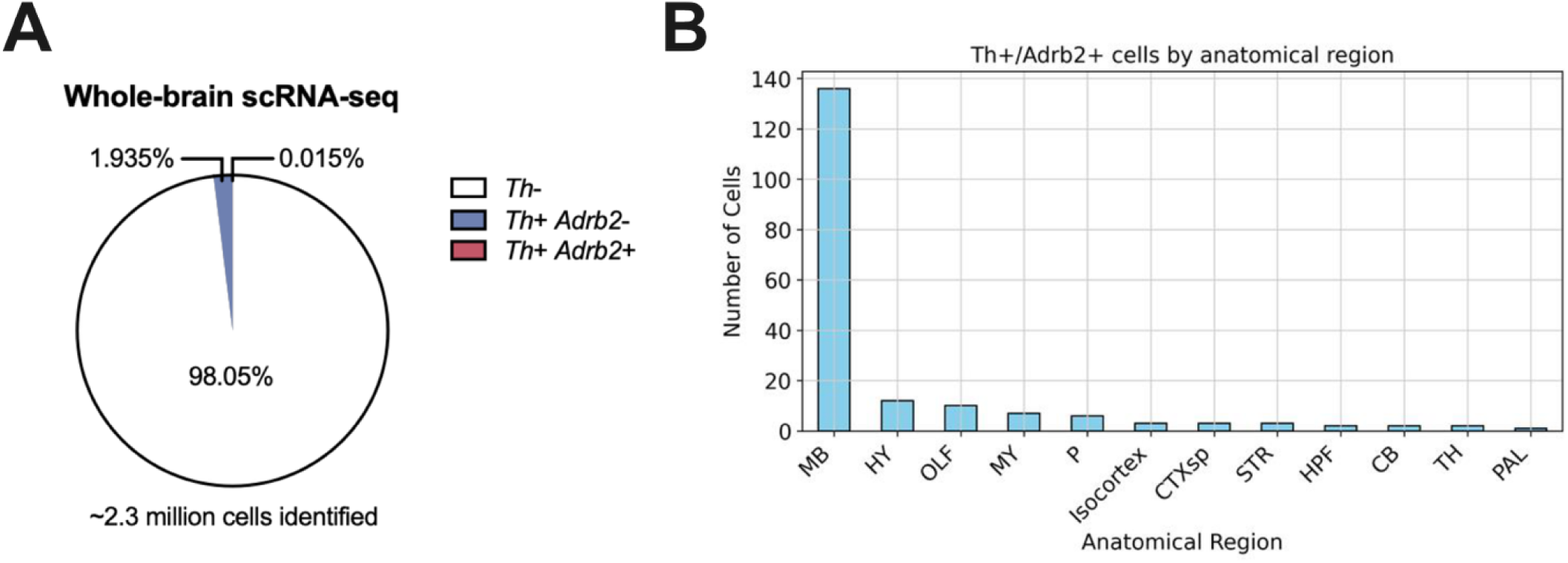
Negligible number of cells in the mouse brain co-expresses *Th* and *Adrb2*. (A) Percentage of cells in whole-brain scRNA-seq dataset positive for *Th* and/or *Adrb2* mRNA. (B) Number of cells co-expressing *Th* and *Adrb2* across brain regions. Midbrain (MB), Hypothalamus (HY), Olfactory areas (OLF), Medulla (MY), Pons (P), Cortical subplate (CTXsp), Striatum (STR), Hippocampal formation (HPF), Cerebellum (CB), Thalamus (TH), Pallidum (PAL).

**Supplementary Table 1.** Details of public scRNA-seq and snRNA-seq datasets for sympathetic ganglia.

| Species | Ganglion Samples | Sequencing Methods | References |
| --- | --- | --- | --- |
| Mus musculus<br>(C57B6 *Wnt1-Cre;Rosa26R<sup>Tomato</sup>*) | Stellate and all thoracic sympathetic ganglia | scRNA-Seq | Furlan et al.<sup>20</sup> |
| Mus musculus<br>(C57B6 *Wnt1-Cre;Rosa26R<sup>Tomato</sup>*) | Stellate and all thoracic sympathetic ganglia | scRNA-Seq | Zeisel et al.<sup>21</sup> |
| Mus musculus<br>(C57B6) | SCG | scRNA-Seq | Mapps et al.<sup>22</sup> |
| Mus musculus<br>(C57B6) | Stellate ganglia<br>(Traced from limb or iBAT) | scRNA-Seq | Lee et al.<sup>23</sup> |
| Mus musculus<br>(C57B6) | Stellate<br>(Traced from heart) | scRNA-Seq | Sharma et al.<sup>24</sup> |
| Mus musculus<br>(C57B6) | SCG and stellate ganglia | snRNA-Seq | Ge et al.<sup>25</sup> |
| Mus musculus<br>(C57B6) | SCG and stellate ganglia | scRNA-Seq | Sarker et al.<sup>26</sup> |
| Mus musculus<br>(C57B6) | SCG | snRNA-Seq | Ziegler et al.<sup>27</sup> |
| Rattus norvegicus | SCG and stellate ganglia | scRNA-Seq | Davis et al.<sup>28</sup> |
| Homo sapiens | Stellate ganglia | snRNA-Seq | Haberman et al.<sup>29</sup> |

